# Integrating cellular graph embeddings with tumor morphological features to predict in-silico spatial transcriptomics from H&E images

**DOI:** 10.1101/2023.10.31.565020

**Authors:** Vignesh Prabhakar, Elisa Warner, Kai Liu

## Abstract

Spatial transcriptomics allows precise RNA abundance measurement at high spatial resolution, linking cellular morphology with gene expression. We present a novel deep learning algorithm predicting local gene expression from histopathology images. Our approach employs a graph isomorphism neural network capturing cell-to-cell interactions in the tumor microenvironment and a Vision Transformer (CTransPath) for obtaining the tumor morphological features. Using a dataset of 30,612 spatially resolved gene expression profiles matched with histopathology images from 23 breast cancer patients, we identify 250 genes, including established breast cancer biomarkers, at a 100 µm resolution. Additionally, we co-train our algorithm on spatial spot-level transcriptomics from 10x Visium breast cancer data along with another variant of our algorithm on TCGA-BRCA bulk RNA Seq. data, yielding mutual benefits and enhancing predictive accuracy on both these datasets. This work enables image-based screening for molecular biomarkers with spatial variation, promising breakthroughs in cancer research and diagnostics.

## Introduction

Spatial transcriptomics, a burgeoning field, has emerged as a powerful tool for unveiling the intricate landscape of gene expression within tissues, shedding light on its profound biological consequences [1]. Traditional transcriptome analyses, whether through bulk sequencing or single-cell sequencing, while informative, fall short in capturing the high-resolution spatial heterogeneity inherent to tissue architecture. The limitations of these approaches become apparent as they often overlook rich spatial information, focusing instead on sampling from various tissue regions [2]. Recent advancements in spatial transcriptomics have addressed this critical gap by introducing innovative techniques that leverage DNA barcodes to distinguish distinct spots within tissues [3]. Each spot can contain tens of cells, and RNA sequencing (RNA-seq) subsequently unravels the expression profiles of these cells. Consequently, hundreds of spots within each tissue section yield gene expression vectors, offering a panoramic view of spatial heterogeneity [4].

Crucially, the spatial transcriptomics approach aligns seamlessly with haematoxylin and eosin (H&E) stained histopathology images routinely available for tissue sections. The pressing challenge is to harmoniously integrate histology images with spatial transcriptomics data to unlock a deeper understanding of gene expression heterogeneity. In this context, machine learning has emerged as a potent ally for comprehensive analysis [5]. While earlier methods relied on hand-crafted features for information extraction and predictions, contemporary approaches harness the capabilities of computer vision algorithms (such as convolutional neural networks and vision transformers) for histopathology image analysis [6]. These techniques have demonstrated remarkable success in tasks such as tumor classification, mutation prediction, and the classification of cancer subtypes [7]. However, their targets of prediction have primarily revolved around slide-level labels rather than fine-grained labels for small cell groups [8].

In this study, we embarked on a pioneering co-training effort, uniting a graph isomorphism model and a vision transformer, and applying them to an extensive dataset encompassing 30,612 spatial transcriptomics spots from 68 breast cancer tissue sections—whole slide images—derived from the 10x Genomics spatial transcriptomics breast cancer dataset [9]. The integration of spatial transcriptomics and histology images through our algorithm is a potent tool for unraveling the intricate gene expression landscape and spatial diversity within breast cancer tissues. Key cell types, including macrophages, epithelial cells, lymphocytes, and neutrophils, hold pivotal roles in this context. Macrophages contribute to the immune response, lymphocytes are central to assessing immune infiltrates, and neutrophils are involved in tumor progression. Epithelial cells form the breast tissue foundation [10-13]. The high-resolution gene expression prediction of our algorithm can enable researchers to discern spatial nuances in these cell types’ functions and interactions, illuminating how they influence one another’s gene expression patterns within the breast tissue microenvironment. This approach provides a comprehensive understanding of breast cancer biology, with potential diagnostic and therapeutic implications.

Our algorithm harnesses the power of both graph isomorphism neural networks, which excel at capturing cell-to-cell interactions from histology images alongside CTranspath - which is a vision transformer pretrained on TCGA whole slide images and can adeptly extract critical tumor morphological features [14][15]. Simultaneously, we harnessed the rich resource of the TCGA BRCA dataset, which provides bulk RNA-seq ground truth at the full whole slide image level across 1977 unique whole slide image samples [16].

Our co-training approach was primarily designed to enhance RNA-seq gene expression predictions for individual spots in the 10x Genomics breast cancer spatial transcriptomics task [9]. This integration yielded notable improvements across various evaluation metrics, including a reduction in Root Mean Squared Error (RMSE), enhanced Area Under the Receiver Operating Characteristic Curve (AUROC), and increased Spearman correlation coefficient scores, especially for predicting breast cancer biomarker gene expressions. Importantly, the advantages of this co-training strategy extended beyond the spatial transcriptomics domain. When we applied the co-trained embeddings to a broader context, utilizing multiple instance learning across entire whole slide images from the TCGA BRCA dataset, we observed remarkable boosts in the Spearman correlation coefficient for breast cancer biomarker gene expression prediction. Additionally, our co-training approach substantially reduced RMSE in downstream regression tasks, resulting in heightened accuracy and precision in predicting gene expressions from bulk RNA-seq data. This comprehensive approach demonstrates the potential of co-training to bridge the gap between high-resolution spatial transcriptomics and bulk RNA-seq, offering profound insights and advancements in breast cancer research.

We demonstrate the robustness of our algorithm by accurately predicting spatial expression patterns in the 10x Genomics breast cancer data. Importantly, it surpasses methods reliant on standard cellular features for expression prediction.

The integration of spot-level spatial transcriptomics and the corresponding histology tissue section for the spots alongside co-training with TCGA-BRCA bulk RNA-Seq data through our model opens new frontiers for dissecting the complex interplay between gene expression and tissue architecture, holding immense promise for cancer research and diagnostic applications. In the subsequent sections, we delve into related work in this space, methodology, results and interpretability of our innovative approach.

## Related Work

The advent of deep learning techniques in the realm of digital pathology analysis has revolutionized clinical practice by offering versatile solutions for a wide array of clinical inquiries, from diagnostic assessments to prognostic predictions. Among the various applications, deep learning models have shown remarkable potential in predicting gene mutations directly from pathology images, yet a comprehensive evaluation of their capacity to extract molecular features from histology slides remains a largely uncharted territory. The HE2RNA paper addresses this gap by introducing HE2RNA, a multifaceted model that seamlessly integrates multiple data modalities [8]. Notably, HE2RNA demonstrates its ability to systematically predict RNA-Seq profiles solely from whole-slide images, eliminating the need for expert annotations. With an interpretable design, HE2RNA offers a virtual spatialization of gene expression patterns, a feature validated through independent staining datasets, such as CD3 and CD20 markers. Moreover, this paper highlights HE2RNA’s transferability across datasets, even those of limited size, to enhance the prediction accuracy of specific molecular phenotypes, showcasing its versatility for clinical applications. This innovative approach is exemplified in the identification of tumors with microsatellite instability, exemplifying its potential to impact clinical diagnosis significantly. While convolutional neural networks (CNNs) have already demonstrated their prowess in enhancing diagnostic capabilities, the marriage of these networks with transcriptomic data offers a promising avenue for deciphering the intricate relationships between molecular signatures and morphological patterns in pathology, ultimately advancing our understanding of disease mechanisms and improving patient care.

The emergence of spatial transcriptomics has significantly advanced our understanding of gene expression patterns within tissues, offering valuable insights into the intricate spatial organization and heterogeneity of gene expression. Conventional transcriptome analyses, whether using bulk or single-cell sequencing, often fail to capture high-resolution spatial heterogeneity, resulting in the loss of critical spatial information. Recent developments in spatial transcriptomics, utilizing DNA barcodes to distinguish distinct spots within tissue sections, have revolutionized our ability to preserve spatial context. This approach allows the recovery of gene expression data for numerous spots within a tissue section, complemented by the availability of corresponding histopathology images. The challenge lies in effectively integrating these histology images with spatial transcriptomics data, a task increasingly tackled by machine learning techniques. While prior methods relied on hand-crafted features, recent advancements, particularly the utilization of convolutional neural networks (CNNs), have demonstrated success in predictive tasks based on histopathology images. However, these efforts typically focused on slide-level labels, overlooking fine-grained cell-level predictions. To bridge this gap, the ST-Net paper introduces a deep learning algorithm, which marries spatial transcriptomics and histology images, enabling high-resolution capture of gene expression heterogeneity [17]. This innovation, trained on a substantial dataset of breast tissue sections from patients with breast cancer, exhibits robust generalization to new samples, surpassing the accuracy of methods reliant on standard cellular features. Moreover, ST-Net excels in predicting substantial variations in expression even within regions classified as tumors by clinicians, thereby highlighting its capability to capture intratumor heterogeneity, a critical facet in the study of cancer biology.

The paper on spatial clustering in transcriptomics presents a timely review of the burgeoning field of spatial transcriptomics, where advancements have enabled the simultaneous profiling of cellular transcriptomes and their spatial context [18]. As these experimental technologies continue to evolve, the need for innovative analytical approaches becomes increasingly evident. The paper highlights the vital role of spatial transcriptomics in offering a new dimension to our understanding of cellular transcriptomes within their native tissue context. It underscores the utility of spatial information in providing novel insights into gene expression patterns. Notably, the paper emphasizes the challenges stemming from the growing volume and complexity of spatial transcriptomics data, necessitating the development of tailored computational methods. While drawing parallels with single-cell RNA sequencing (scRNA-seq) data analysis, the paper advocates for novel approaches that harness the unique spatial dimension offered by spatial transcriptomics. It highlights the synergistic potential of integrating spatial transcriptomics with scRNA-seq data, demonstrating improvements in various research areas. The review categorizes the computational methods employed in spatial transcriptomics data analysis, offering a comprehensive overview of the current landscape, and concludes by outlining future challenges and opportunities in this rapidly evolving field. Overall, this paper serves as a valuable resource to inspire future method development and advancements in spatial transcriptomics data analysis.

The SpaCell paper introduces a groundbreaking approach to spatial transcriptomics (ST) data analysis, addressing a crucial gap in current methodologies [19]. While ST technology offers the unique advantage of mapping gene expression to specific spatial locations within intact tissue, it has largely underutilized the valuable information embedded in pixel intensity data from tissue images. The paper presents SpaCell, a user-friendly deep learning software that seamlessly integrates millions of pixel intensity values with gene expression measurements, thereby bridging the gap between molecular data and tissue morphology. Unlike traditional analysis pipelines that focus solely on gene expression, SpaCell harnesses both data types to construct neural network models for cell-type identification and disease-stage classification. This novel integration approach outperforms methods using gene-count or imaging data in isolation, providing high-resolution and accurate predictions. The paper underscores the potential of combining genomics and imaging data for innovative machine learning applications in characterizing tissue morphology and disease staging, a promising avenue in the field of spatial transcriptomics.

The paper on spatial gene expression analysis in HER2-positive breast tumors employs Spatial Transcriptomics (ST) technology to uncover intricate cellular interactions and spatial dimensions within the tumor microenvironment [20]. While transcriptomic studies have greatly enhanced our understanding of cancer, the loss of spatial information in traditional sequencing strategies hinders the exploration of cell-to-cell interactions critical for disease progression. HER2-positive breast cancer serves as the context for this investigation, as it represents a subtype demanding advanced treatment strategies. The study leverages ST to unravel the spatially resolved gene expression profiles in 36 samples from eight HER2-positive patients. Through expression-based clustering and integration with single-cell data, the paper identifies shared gene expression signatures, high-resolution cell state colocalization patterns revealing an interferon-associated macrophage T-cell interaction, and introduces a method for detecting potential tertiary lymphoid-like structures within the tumor. This comprehensive analysis bridges the gap between molecular data and tissue morphology, shedding light on intricate cellular relationships in HER2-positive breast tumors and offering insights into potential treatment strategies and immune system involvement in this context.

Spatial transcriptomics has emerged as a powerful tool to analyze gene expression patterns within the context of tissue organization, but parsing complex datasets remains challenging. Existing methods for identifying spatial gene expression patterns often rely on statistical hypothesis testing, motivating the exploration of alternative approaches. In this study, the authors propose a novel strategy that simulates transcript diffusion to extract genes exhibiting distinct spatial patterns. By measuring the time required for transcripts to converge within a simulated spatial domain, they rank gene expression profiles based on their spatial organization. This unorthodox approach, implemented in an open-source tool named “sepal,” offers a complementary perspective to existing methods and is less influenced by gene expression levels [21]. The method’s potential to discern structured spatial patterns in transcriptomic data is demonstrated on both array-based and unstructured spatial transcriptomics platforms, showcasing its versatility in large-scale spatial data analysis. This innovative approach provides a valuable addition to the toolkit for exploring spatial gene expression patterns.

Spatially resolved transcriptomics (SRT) has transformed our ability to explore gene expression patterns in the context of tissue microenvironments. To harness the wealth of information provided by SRT data, the SpaGCN method is introduced, leveraging graph convolutional networks (GCNs) to integrate gene expression, spatial location, and histology [22]. Unlike traditional clustering methods, SpaGCN identifies spatial domains characterized by coherent gene expression and histological features, accounting for the spatial dependency of gene expression in a flexible manner that is compatible with various SRT platforms. Additionally, SpaGCN jointly addresses the identification of spatially variable genes (SVGs) within these domains, ensuring that the detected genes exhibit genuine spatial expression patterns. This holistic approach provides a comprehensive understanding of spatial gradients in gene expression within tissues, making SpaGCN a versatile tool for analyzing diverse SRT datasets.

The GCNG (Graph Convolutional Neural network approach for Genes) method addresses the challenge of inferring gene interactions from high-throughput spatial transcriptomics data, particularly focusing on extracellular interactions between genes in different cells [23]. While previous methods primarily concentrated on intra-cellular interactions, the advent of spatial transcriptomics has opened new avenues for studying both intra- and inter-cellular interactions within tissues. GCNG leverages graph convolutional neural networks (GCNs) to encode spatial information as a graph and integrate it with gene expression data through supervised training. This approach enhances the identification of gene interactions between cells and can even propose novel pairs of interacting extracellular genes. Moreover, GCNG’s output can be valuable for downstream functional assignments, offering a promising tool for analyzing spatial transcriptomics datasets and uncovering complex cellular interactions.

The STlearn paper presents groundbreaking advancements in the field of spatial transcriptomics (ST), a technology that combines gene expression data, spatial location information, and tissue morphology to enhance our understanding of cellular interactions within intact tissues [24]. ST data offers tremendous potential for deciphering the biology of cell types in their native tissue context. The paper introduces innovative analysis approaches implemented in the stLearn software to leverage all three data types effectively. These approaches address three key research areas: identifying cell types, reconstructing cell type evolution within tissues, and studying cell-cell interactions. By integrating spatial information and cellular morphology with gene expression measurements, the methods developed in this study significantly enhance the sensitivity and accuracy of cell type identification, cell trajectory reconstruction, and the detection of cell-cell interactions within tissues. The stLearn software provides a comprehensive toolkit for processing and analyzing ST data, offering researchers a powerful platform to gain insights into complex biological processes within both healthy and diseased tissues.

The spatialLIBD paper presents a comprehensive exploration of gene expression patterns within the six-layered human dorsolateral prefrontal cortex (DLPFC) using the 10x Genomics Visium platform for spatial transcriptomics [25]. This groundbreaking study leverages spatial transcriptomics to gain insights into the spatial organization of gene expression within the human brain, particularly focusing on the distinct gene expression patterns across cortical layers. By overlaying the obtained laminar expression signatures with large-scale single nucleus RNA-sequencing data, the authors enhance spatial annotation of expression-driven clusters. Furthermore, they integrate gene sets related to neuropsychiatric disorders, revealing differential layer-enriched expression of genes associated with conditions such as schizophrenia and autism spectrum disorder, underscoring the clinical relevance of spatially defined expression. Additionally, the paper introduces a data-driven framework for unsupervised clustering in spatial transcriptomics data, a valuable tool applicable to various tissues and brain regions where morphological architecture is not as clearly defined as in the cortex. The authors make their data and analysis tools available to the scientific community, offering a valuable resource for advancing neuroscience research and spatial transcriptomics investigations in the human brain.

This paper introduces a groundbreaking approach called expression-morphology (EMO) analysis in breast cancer, leveraging deep convolutional neural networks (CNNs) to predict mRNA expression patterns and molecular markers from routine histopathology whole slide images (WSIs) [26]. This innovative method aims to harness the rich morphological information present in hematoxylin and eosin (H&E) stained tissue sections to infer the underlying molecular phenotypes of tumors. The study covers a transcriptome-wide analysis with gene-specific CNN models, achieving significant associations between predicted expressions and RNA sequencing estimates for thousands of genes. Importantly, EMO analysis provides insights into intratumor heterogeneity by predicting spatial expression patterns, which are subsequently validated using spatial transcriptomics profiling. Furthermore, the application of EMO for predicting a clinically relevant proliferation score underscores its potential for precision medicine in breast cancer. This comprehensive approach not only represents a cost-efficient and scalable means of predicting tumor molecular phenotypes but also offers insights into intratumor variability and clinical applications, making it a valuable tool for cancer research and diagnostics.

This paper presents a novel deep learning-based approach for predicting RNA-sequence expression (RNA-seq) from Hematoxylin and Eosin whole-slide images (H&E WSI) in head and neck cancer patients [27]. The study addresses the challenge of efficiently leveraging histopathological images to infer molecular profiles, which can be invaluable for targeted cancer therapies. Unlike conventional methods that predict RNA-seq at a patch-by-patch level, potentially losing crucial spatial-contextual relationships, the proposed framework employs a neural image compressor to capture these relationships and generate a compressed representation of the whole-slide image. A customized deep-learning regressor then predicts RNA-seq by learning both global and local features. Testing on a TCGA-HNSC dataset demonstrated superior performance, achieving higher mean correlations and improved predictions for several oncogenes compared to state-of-the-art baseline methods. The interpretability of the results, including pathway analysis and activation maps, adds to the clinical relevance of this approach, potentially enabling the discovery of genetic biomarkers directly from histopathology images and streamlining patient pre-screening before genetic testing, thereby saving time and costs.

This paper addresses the crucial problem of linking genomic data to phenotypes using histopathological images, offering a unique approach to bridge the gap between genes and cellular tissue organization [28]. While genomics has made significant strides, histopathological images remain essential for precise oncological diagnoses, underscoring the need for integration. The study leverages deep learning techniques to extract visual features from histological images, eliminating the subjectivity associated with human histopathologists’ evaluations. These objective visual features are then correlated with gene expression profiles, providing a more quantifiable relationship between transcriptomes and phenotypes. The research, focused on normal tissues from the Genotype-Tissue Expression (GTEx) dataset, offers a novel path toward understanding the links between genes, their expression, and histological phenotypes, with potential applications in improving cancer prognosis and treatment strategies. The challenges of working with large-scale whole-slide images and the diverse array of deep learning architectures in the field are also discussed, highlighting the ongoing developments and promising prospects in the realm of digital histopathology.

This paper addresses the challenge of distinguishing between adenocarcinoma, squamous cell carcinoma, and healthy lung tissue, the two most prevalent lung cancer types, using an integrative classification approach [29]. By combining histology and RNA-Seq data, the authors propose a late fusion classification model that significantly improves diagnostic accuracy compared to using each data source independently. The model demonstrates a remarkable reduction in the diagnosis error rate, achieving a mean F1-Score of 95.19% and a mean AUC of 0.991. These results underscore the potential of leveraging multiple sources of biological data to enhance the accuracy of cancer subtype diagnosis. Importantly, the late fusion methodology can be adapted to various cancer types or diseases characterized by diverse information sources, offering a promising approach for improving clinical decision support systems and advancing personalized medicine.

## Methods

### Data Acquisition and Preprocessing

To validate our approach using spatial transcriptomic data, we obtained publicly accessible breast cancer tissue slides and spatial transcriptomic datasets processed using the Spatial Transcriptomics method [30]. This dataset encompassed breast tissue samples from 23 patients, providing spot coordinates for the spatial transcriptomics assay on each slide, along with their corresponding RNA-seq expression values. These slides included multiple replicates of various breast cancer subtypes, including luminal a, luminal b, triple-negative, and HER2-positive, resulting in a total of 68 samples. Prior to data acquisition, the tissue images underwent scanning, freezing, and embedding in an OCT compound. Subsequently, they were sequenced using a specialized assay that incorporated positional molecular barcodes, referred to as “spatial barcodes,” in addition to standard RNA-seq reagents [17][31]. These unique barcodes enabled precise spatial mapping of gene expression. To prepare the histology input for our model, we extracted image tiles sized at 224x224 pixels from each slide, with each image centered around the pixel coordinate of an assay spot, representing a 100μm region around the spot. For our reference gene analysis, we obtained a total of 26,949 raw sequence counts for each spot, which were then subjected to log-normalization. All analyses were carried out using Python 3.10.

### Train-test-validation split

In our experimental design, we undertook a meticulous train-test split procedure to rigorously assess the performance of our models. For model selection, we opted for a leave-one-patient-out cross-validation approach, given the availability of 23 distinct patients’ profiles and a corresponding set of approximately 69 H&E images (roughly three H&E images per patient profile). This cross-validation strategy allowed us to thoroughly evaluate the models’ generalization capabilities and robustness across different patient profiles. Additionally, to provide some final insights and results, we conducted a fresh 80-20 train-test split on the dataset, generating sample predictions. This approach enabled us to assess the models’ performance on a previously unseen dataset, offering valuable insights into their predictive power and applicability beyond the training data.

### Algorithm Design and Training

As shown in Figure 1, the first step involves constructing a cellular graph from H&E tiles where nodes represent individual cells segmented and extracted from the tiles using Hovernet and edges signify spatial proximity based on K-nearest neighbors. Next, we employ Graph Isomorphism Network (GIN) embeddings to generate node-level representations to capture critical cellular features and interactions. Simultaneously, we leverage CTransPath embeddings to capture tumor morphology features from the same set of H&E tiles. These embeddings are then concatenated together to create a fused, comprehensive embedding. Subsequently, a multi-output regression model is trained to predict spatial transcriptomic profiles, using these joint embeddings as features and normalized gene expression values as ground truth. This integrated framework not only enhances prediction accuracy but also facilitates interpretability. Furthermore, we perform co-training with TCGA BRCA bulk RNA-Seq data to boost predictive power and generalize findings across different datasets, ensuring a holistic understanding of the intricate gene expression landscape within breast cancer tissues.

**Figure 1.**
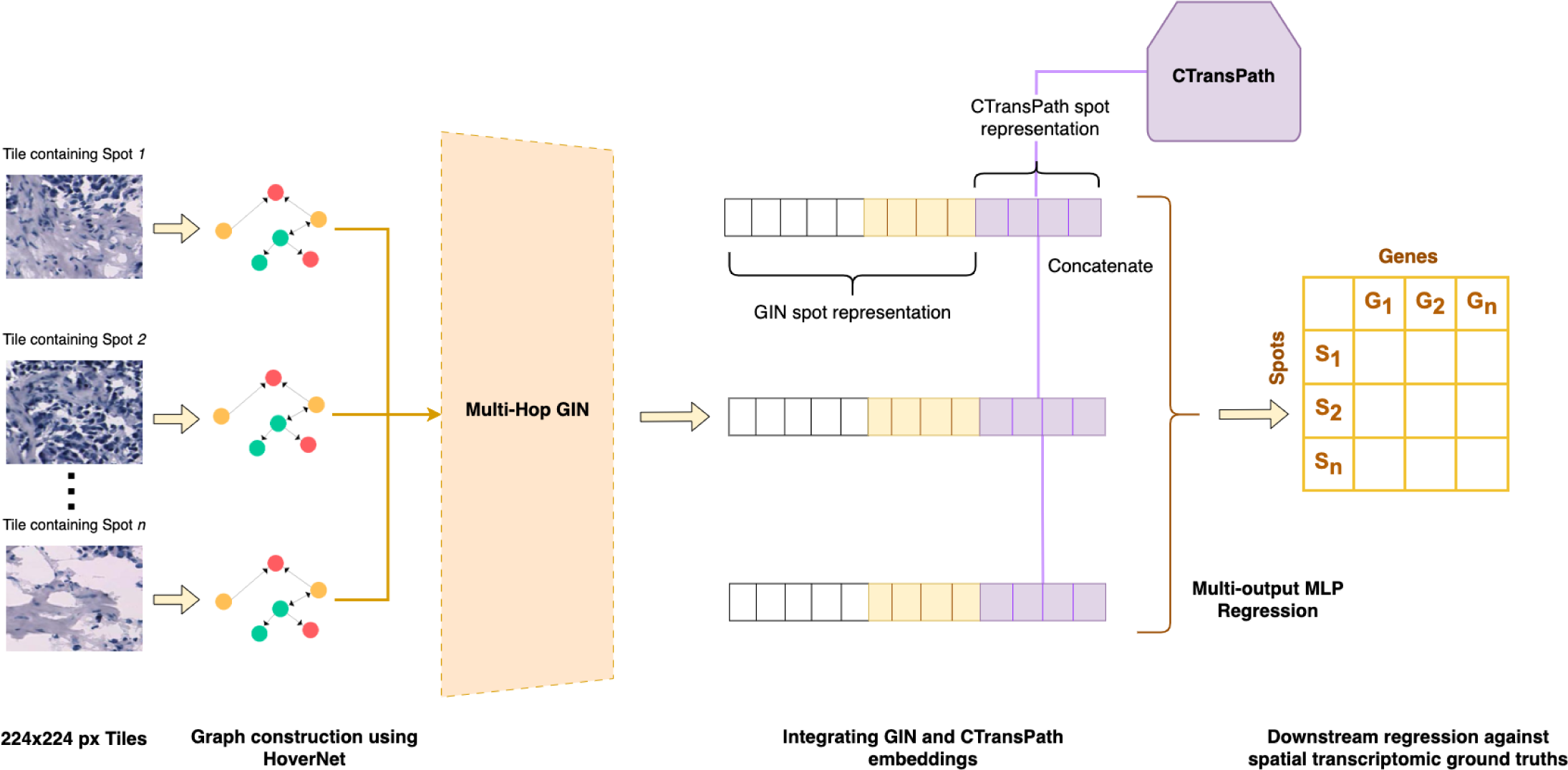
End to end architecture of our model that integrates graph embeddings (representing cell-cell interactions at a given tile/spot) with vision embeddings (representing tumor morphological features at a given tile/spot) in order to generate spot representations for predicting the in-silico spatial transcriptomics of the breast cancer tissues sample using downstream multi-output MLP regression.

### Cell Segmentation and Classification

Hover-Net is a state-of-the-art deep learning framework tailored for cell analysis in H&E-stained histopathological images [32]. It leverages the power of convolutional neural networks (CNNs) and demonstrates remarkable accuracy in segmenting individual cells and classifying them into distinct cell types. Hover-Net excels in semantic segmentation, accurately delineating the boundaries of individual cells within H&E images. This is achieved through a combination of convolutional and deconvolutional layers. It goes beyond segmentation and classifies each segmented cell into predefined categories, such as epithelial, lymphocyte, macrophages or neutrophils. The network effectively captures contextual information, enabling it to differentiate between cells with subtle morphological variations. Hover-Net is designed for efficiency, ensuring fast and reliable results, making it suitable for real-time or large-scale applications.

### Cellular Graph Construction

Once we have accurately identified and classified individual cells, the next crucial step is to construct a meaningful representation of their spatial relationships within the tissue sample. To achieve this, we utilize a graph-based approach as shown in Figure 2, where nodes in the graph correspond to the individual cells identified by Hover-Net.

**Figure 2.**
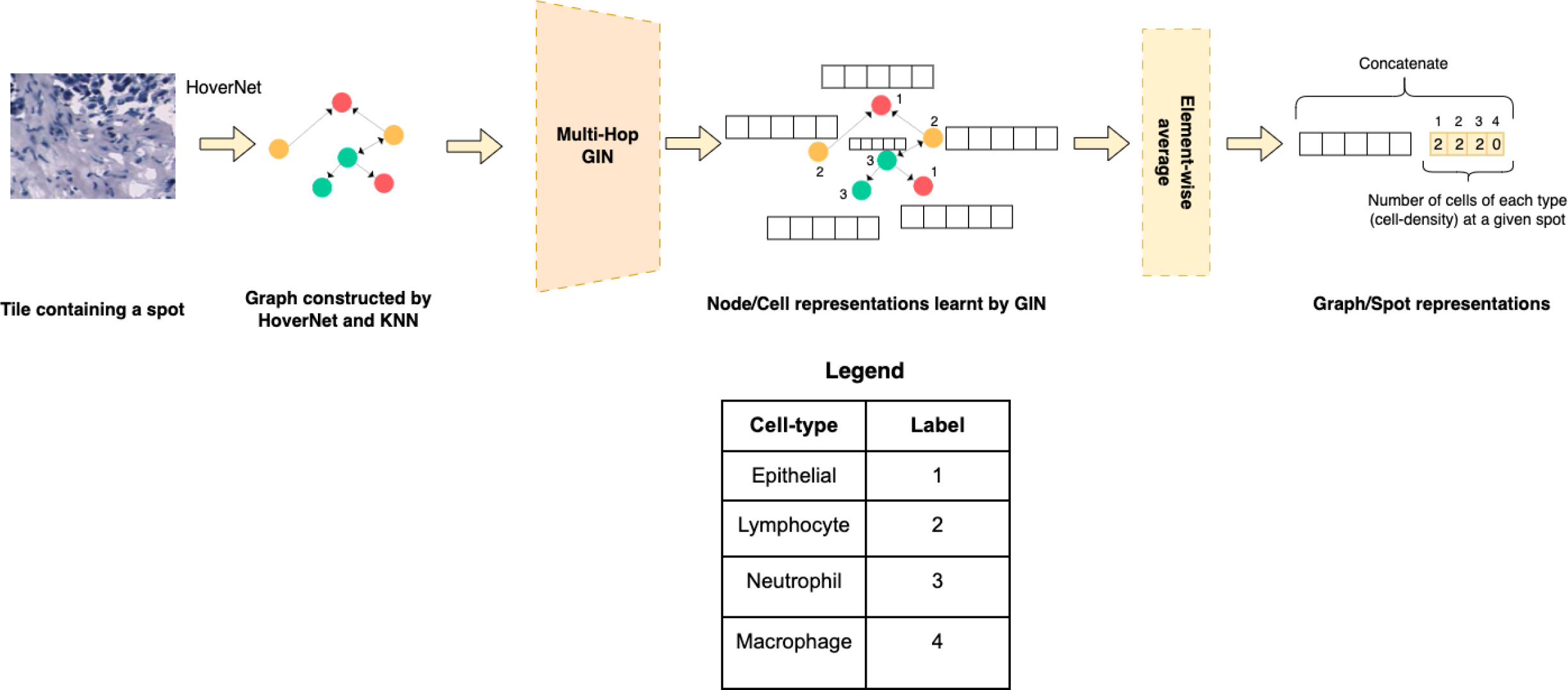
The steps involved in cellular graph construction for a given tile/spot followed by the generation of node/cell representations for each cell which in turn help generate spot representations using simple aggregation methods such as an element-wise average. The resultant graph/spot representation from this sequence of steps is used alongside vision embeddings for the same tile/spot during downstream regression. The cells captured as a part of the graph construction procedure include Epithelial cells, Lymphocytes, Neutrophils and Macrophages as shown in the legend.

Each cell is treated as a node in the graph, and we establish edges (connections) between these nodes based on spatial proximity. Specifically, we employ a K-nearest neighbors (K = 5) algorithm to identify the five closest neighboring cells for each cell in the image [33]. These nearest neighbors represent the immediate spatial context of the focal cell. Edges are then established between the focal cell and its five neighbors, creating a network that encapsulates the spatial relationships between neighboring cells.

This graph construction approach allows us to capture the intricate cellular interactions and arrangements within the tissue sample, providing valuable insights into the spatial organization of cells. The resulting graph becomes a powerful representation for further analysis, facilitating tasks such as studying cellular neighborhoods, identifying potential clusters or patterns, and aiding in the interpretation of complex tissue structures. This graph-based spatial analysis, coupled with the precision of Hover-Net, enhances our ability to extract meaningful information from the H&E images and contributes significantly to our research objectives.

### Generating Graph Isomorphism Network (GIN) embeddings for each tile encompassing a cellular graph

In our research framework, we take advantage of Graph Isomorphism Networks (GIN) to extract informative node embeddings from the graph constructed based on cell spatial relationships within histopathological images [14]. Each node in this graph represents an individual cell, and the edges denote spatial proximity between cells, elucidating their contextual interactions. Leveraging GIN, we train a deep learning model to generate node embeddings for each cell. GIN excels at capturing local and global graph structures, making it a potent choice for our task.

Subsequently, we extend this node-wise representation to obtain graph embeddings for each larger entity, known as a “spot,” which encompasses multiple cells within a specific region delineated by the image tile. To achieve this, we adopt an aggregation approach, where we average the node embeddings of all constituent cells within a given spot. This aggregation step results in comprehensive graph embeddings that encapsulate the spatial organization and relationships among cells within the respective spots.

These graph embeddings serve as crucial features that encode the spatial transcriptomic ground truths for the corresponding image tiles. By employing GIN and this graph-based embedding strategy, our research aims to unveil the intricate interplay between cell spatial arrangements and molecular expression patterns, thus advancing our understanding of tissue biology and disease mechanisms.

### Generating CTransPath features for each tile in the WSI

In our research, we leverage the powerful CTransPath model to extract vital visual features from histopathological image tiles, particularly focusing on the H&E spots enclosed within these tiles [15]. The CTransPath model is pre-trained using a unique self-supervised learning strategy known as semantically-relevant contrastive learning (SRCL). This strategy, unlike traditional contrastive learning, enhances the diversity of positive instances by aligning multiple instances that share similar visual concepts. This augmentation of positive pairs leads to the generation of highly informative representations, which are specifically tailored to capture intricate histopathological characteristics.

Comparatively, this approach with CTransPath differs from the GIN model previously mentioned. While GIN concentrates on constructing a graph-based representation of cells within the spot and averages their node embeddings to obtain a comprehensive graph embedding for the entire spot, CTransPath operates at a different level. It hones in on the pixel-level visual details of the histopathological image, extracting features that encapsulate the fine-grained visual attributes of the enclosed H&E spot. These features hold rich information about the staining patterns, cell structures, and other histological nuances that are distinct from the graph-based spatial relationships captured by GIN. By employing CTransPath’s SRCL-pretrained model, we aim to provide a complementary perspective in our research, facilitating a holistic understanding of the complex interplay between spatial organization and visual characteristics within histopathological images.

### Integration of CTransPath embeddings with cellular graph embeddings

We take a comprehensive approach to harness the strengths of both the GIN (Graph Isomorphism Network) and CTransPath models to enrich our insights into histopathological images. Following the acquisition of distinct embeddings from these models for each H&E spot within the image tiles, we recognize the significance of integrating their unique perspectives. To accomplish this, we simply concatenate the GIN and CTransPath embedding vectors in order to create a unified embedding representation for each individual spot. This fusion strategy amalgamates the spatial insights captured by GIN, reflecting cellular interactions and connectivity within the spot, with the fine-grained visual features extracted by CTransPath, representing the pixel-level histopathological characteristics [34].

This integrative approach amplifies the richness of the embeddings, bolstering our research with a multi-dimensional representation of histopathological data.

### Downstream Multi-output MLP Regression

In our study, we employed a sophisticated multi-output regression framework to establish a direct link between the joint embeddings derived from the integrated GIN and CTransPath models and the spatial transcriptomic normalized gene expression values. By treating the normalized gene expression values as our ground truths, we aimed to leverage the rich information encoded in the joint embeddings to predict the gene expression profiles accurately. This approach enables us to capture the intricate relationships between the histopathological characteristics represented in the embeddings and the corresponding molecular signatures expressed within the tissue. The multi-output regression model allows us to simultaneously predict multiple genes’ expression levels, reflecting the comprehensive molecular landscape of the histopathological samples [35].

### Co-training 10x Visium breast cancer ST data with TCGA BRCA bulk RNA Seq data

In addition to training the model on the 10x Visium data and its corresponding spatial transcriptomics ground truths, we implemented a co-training strategy that involved another variant of the model as shown in Figure 3. This secondary model was designed to extract embeddings from TCGA BRCA H&E images and utilized whole slide-level bulk RNA Seq data as ground truths. By co-training these two models simultaneously, we aimed to enhance their predictive capabilities by modifying their latent representations for every iteration of co-training. Notably, this co-training approach led to significant improvements in the model’s performance. It not only enhanced the accuracy of predicting spot-level spatial transcriptomics from the 10x Visium data but also resulted in improved predictive power when it came to regressing against the whole slide image-level bulk RNA Seq ground truths in TCGA-BRCA. This joint training strategy ultimately strengthened the model’s ability to make precise predictions across various scales, contributing to its overall effectiveness in bridging histopathology and genomics data.

**Figure 3.**
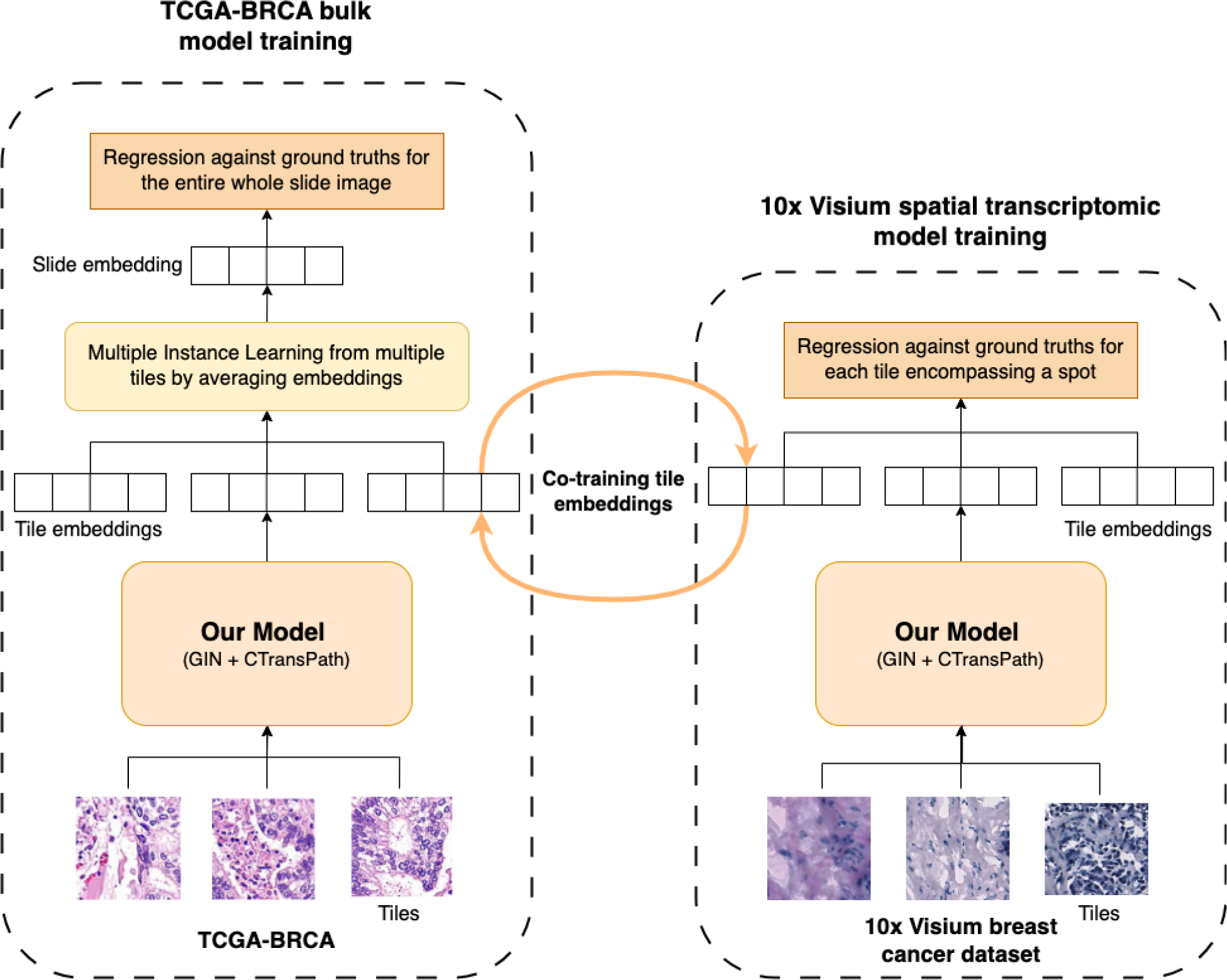
In a symbiotic approach, we co-train the integrated tile embeddings produced by our model on both the TCGA-BRCA and 10x Visium breast cancer datasets, harnessing their distinct latent representations to predict the spatial transcriptomics of breast cancer tissue samples within both these datasets.

## Evaluation and Results

### Evaluation metrics

For evaluating the spatial transcriptomic ground truth prediction task using the multioutput regressor in the context of 10x Visium data with spot-level ground truths, we employed a leave-one-patient-out cross-validation approach. This method involved systematically excluding one patient’s data from the training set while utilizing the remaining patients’ data for model training. The excluded patient’s data were then used as the test set. We computed the Root Mean Squared Error (RMSE) between the predicted spatial transcriptomic values and the actual ground truth values for each spot within the test patient’s data. This process was repeated for each patient in the dataset, and the RMSE values were averaged to obtain an overall assessment of prediction accuracy. Additionally, we utilized the Spearman correlation coefficient to evaluate the relationship between the predicted gene expression values and the actual values for specific biomarker genes, offering insights into the model’s ability to capture the expression patterns of these key genes.

Similarly, for evaluating the predictions in the context of TCGA BRCA data with bulk RNA Seq data, we employed the RMSE metric. Here, we assessed the model’s performance by calculating the RMSE between the predicted bulk RNA Seq values and the corresponding actual ground truth values, providing a quantitative measure of predictive accuracy for this task.

The Spearman correlation coefficient was also utilized in both scenarios, specifically for comparing the predicted gene expression values generated by the model against the actual expression values for a set of biomarker genes. This correlation coefficient served as a valuable indicator of the model’s ability to capture the relative trends and associations in gene expression levels, particularly for genes of interest that hold significance in the context of the study.

Evaluating the advantages of co-training on the TCGA BRCA dataset, we observed a notable improvement in predictive performance. As shown in Figure 4a; prior to co-training, the correlation scores for the predicted value of each gene expression vector versus the actual gene expression vector demonstrated a moderate alignment. However, post co-training, we witnessed a significant boost in these correlation scores as shown in Figure 4b. On average, the scores surged by approximately 4 points, signifying a substantial enhancement in prediction accuracy. This increment highlights the substantial benefits of incorporating co-training into our framework, reinforcing its effectiveness in aligning gene expression profiles and enhancing our understanding of the complex spatial transcriptomics within breast cancer tissues.

**Figure 4a.**
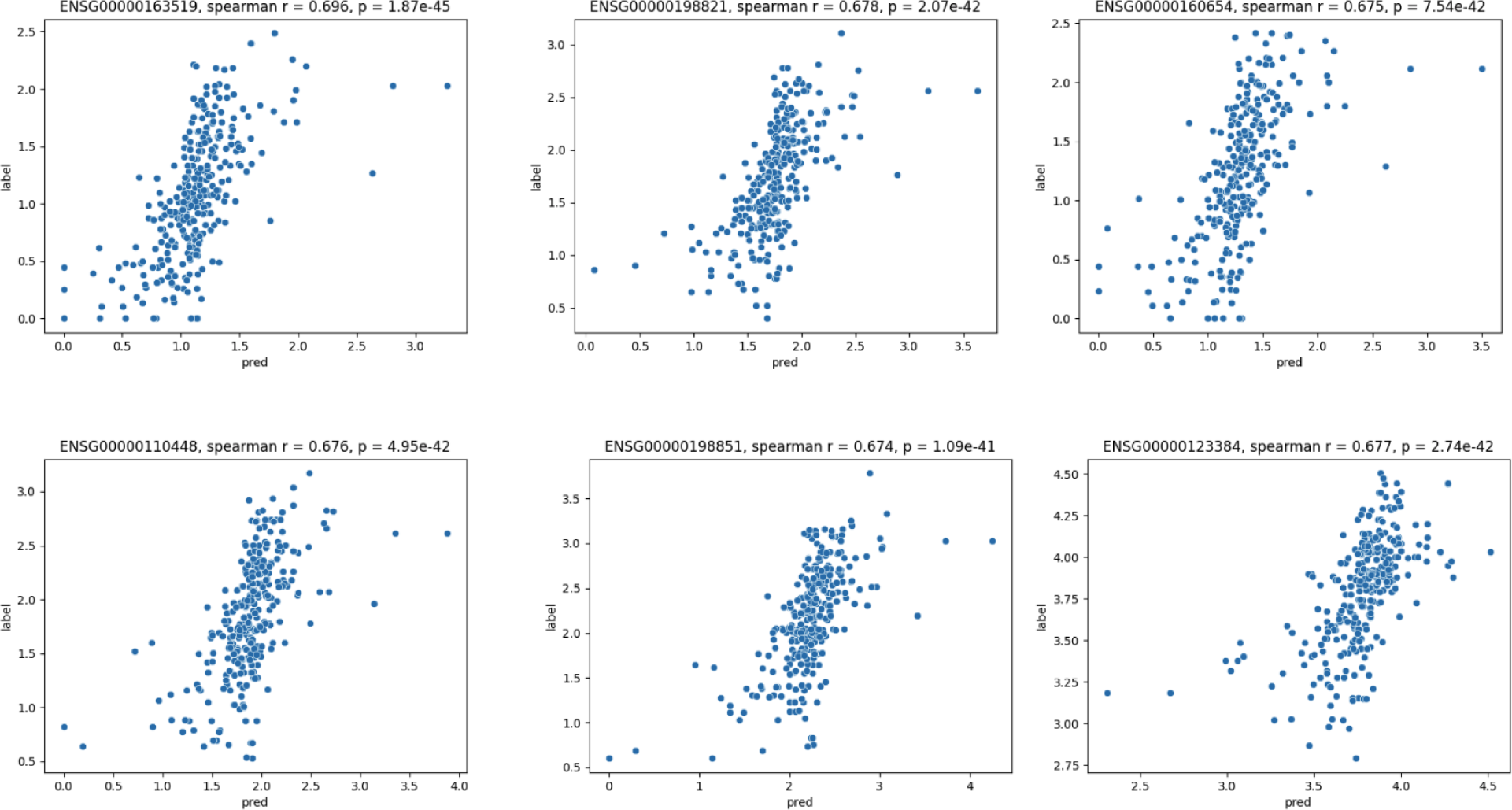
The spearman correlation coefficient plots between the actual gene expression values vs the predicted gene expression values in TCGA-BRCA breast cancer tissue samples for a set of 6 genes prior to co-training with the 10x Visium breast cancer dataset (TRAT1 - ENSG00000163519, CD247 - ENSG00000198821, CD3G – ENSG00000160654, CD5 - ENSG00000110448, CD3E - ENSG00000110448, LRP1 - ENSG00000123384).

**Figure 4b.**
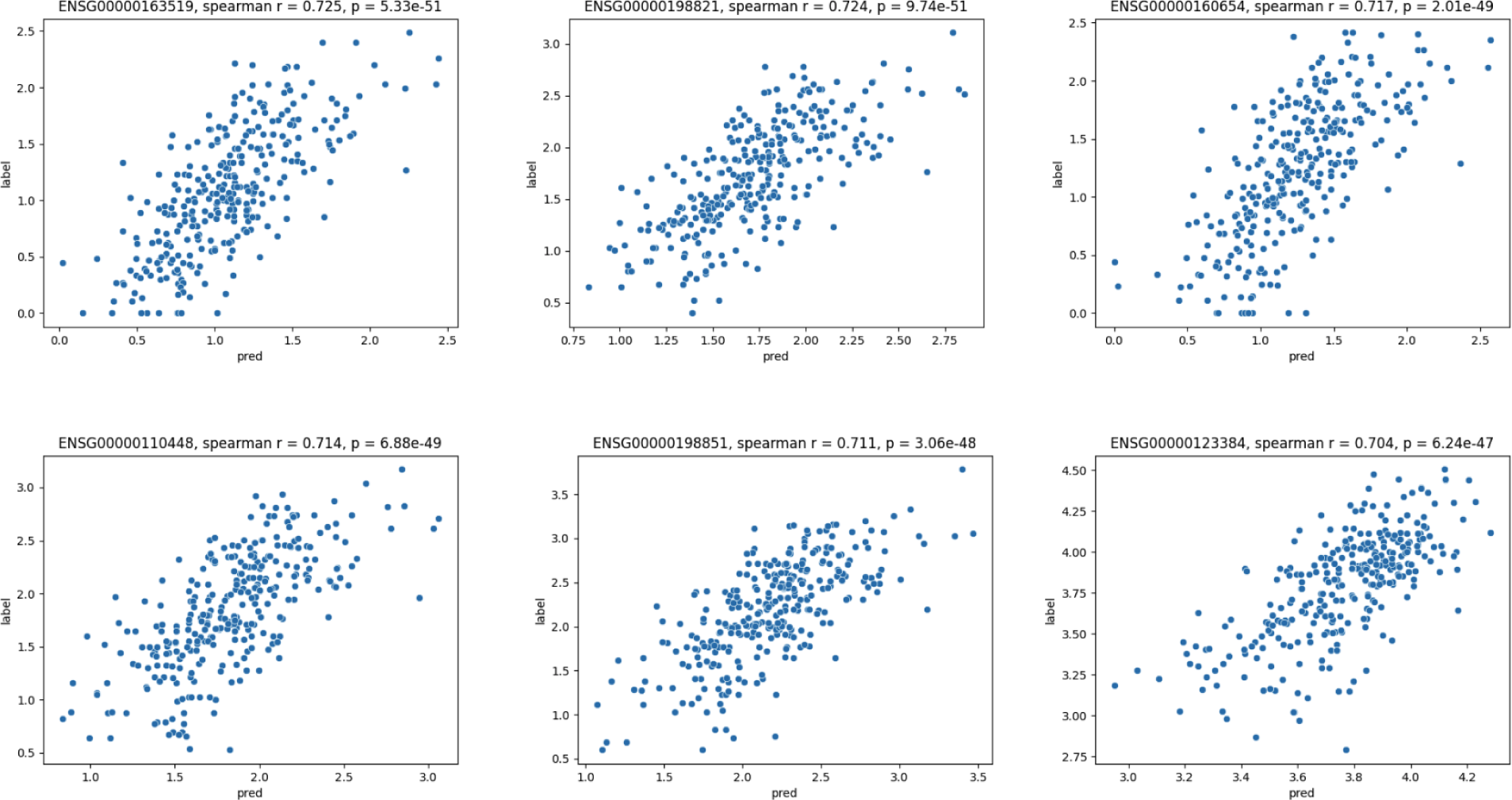
The spearman correlation coefficient plots between the actual gene expression values vs the predicted gene expression values in TCGA-BRCA breast cancer tissue samples for a set of 6 genes after co-training with the 10x Visium breast cancer dataset (TRAT1 - ENSG00000163519, CD247 - ENSG00000198821, CD3G - ENSG00000160654, CD5 - ENSG00000110448, CD3E - ENSG00000110448, LRP1 - ENSG00000123384).

Subsequent to the co-training process, we evaluated the predictive power of our model by measuring the Spearman correlation coefficients between predicted gene expression values, derived from our co-trained model embeddings, and the actual gene expression values in the 10x Visium breast cancer dataset. Our analysis, as illustrated in Figure 5(b), revealed a consistent Spearman correlation coefficient of approximately 0.35 across all the genes encompassed in our selected geneset. This signifies that our co-trained model consistently achieved strong predictive performance, capturing the intricate relationships between gene expressions within the spatial transcriptomic context of breast cancer.

**Figure 5.**
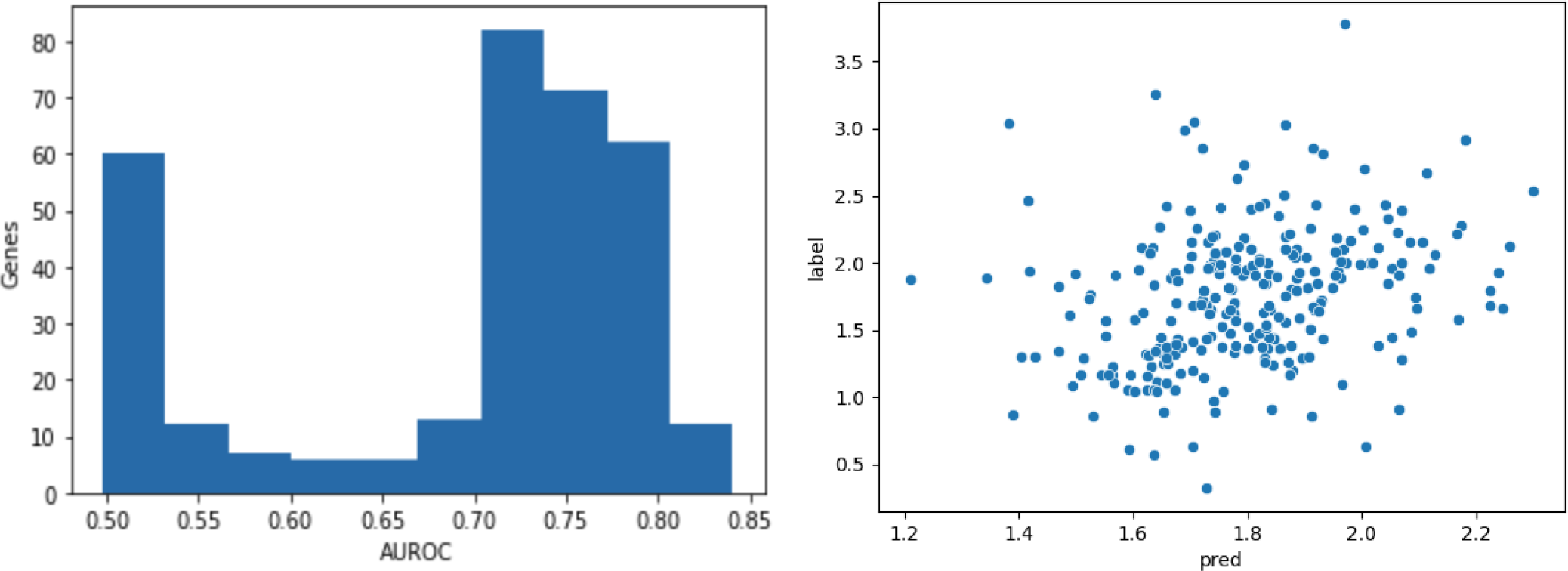
(a) Histogram of AUROC for spatial gene expression predictions using the co-trained embeddings on the test split of the 10x Visium breast cancer dataset. (b) Comparison of spearman correlation coefficients between predicted gene expression values from our co-trained model embeddings and actual values in the 10x Visium breast cancer dataset.

Additionally we also employed the Area Under the Receiver Operating Characteristic Curve (AUROC) metric to assess the model’s ability to distinguish between different levels or classes of predictions. Although AUROC is traditionally used for binary classification tasks, it can also be adapted for regression tasks by treating them as classification problems with a defined threshold. We first divided the range of predicted values into multiple discrete classes or bins. This discretization process involved selecting appropriate thresholds to define the classes. With the thresholds in place, we transformed the regression task into a binary classification problem. Each data point was assigned to one of the classes based on whether its predicted value fell below or above the chosen threshold. Once the data points were categorized into classes, we computed the AUROC as usual. The AUROC measures the ability of the model to correctly rank the samples in terms of their predicted values. It assesses the model’s capacity to distinguish between classes and make accurate predictions relative to a random classifier. The resulting AUROC value provided an indication of the model’s performance in separating the different expression level categories, which is valuable for assessing the accuracy and granularity of spatial transcriptomic predictions.

The AUROC scores showcasing the model’s performance in gene expression vector prediction tasks on 10x Visium breast cancer dataset were also obtained following the co-training process. Our results indicate that after the co-training phase, the model consistently achieved AUROC scores exceeding 0.5 for all genes analyzed. Impressively, over 70% of the genes within the gene set exhibited AUROC scores surpassing the 0.7 threshold, underscoring the model’s heightened predictive accuracy and reliability across a diverse set of genes relevant to breast cancer research. These observations, as visually depicted in Figure 5(a), reinforce the effectiveness of our co-training approach in enhancing predictive capabilities.

### Enrichment of known biomarker genes for breast cancer

In our evaluation of spatial transcriptomic prediction, we conducted an essential analysis to gauge the enrichment of five well-established breast cancer biomarker genes: GNAS, ACTG1, FASN, DDX5, and XBP1 [17]. These genes, recognized for their significance in breast cancer, serve as crucial indicators of disease characteristics and progression. By assessing their presence and expression patterns within the predicted results as shown in Table 1, we gain valuable insights into the model’s accuracy and its ability to identify key biomarkers associated with breast cancer. This analysis adds an important dimension to our research, reinforcing the relevance of our predictive model in the context of breast cancer diagnosis and prognosis. The objective of Table 1 is to showcase the predicted gene expression levels of these biomarker genes across all 30,612 tumor spots within the dataset and thereby offer insights into the enrichment patterns of these five known biomarker genes within the spatial context of 10x Visium breast cancer tissue samples. To achieve this, we utilized our co-trained model embeddings to calculate gene expression predictions for each spot. We then identified the top 10 highly expressed genes across all the tumor spots and measured the number of spots where the biomarker genes are one among the top 10 highly expressed genes.

**Table 1.**
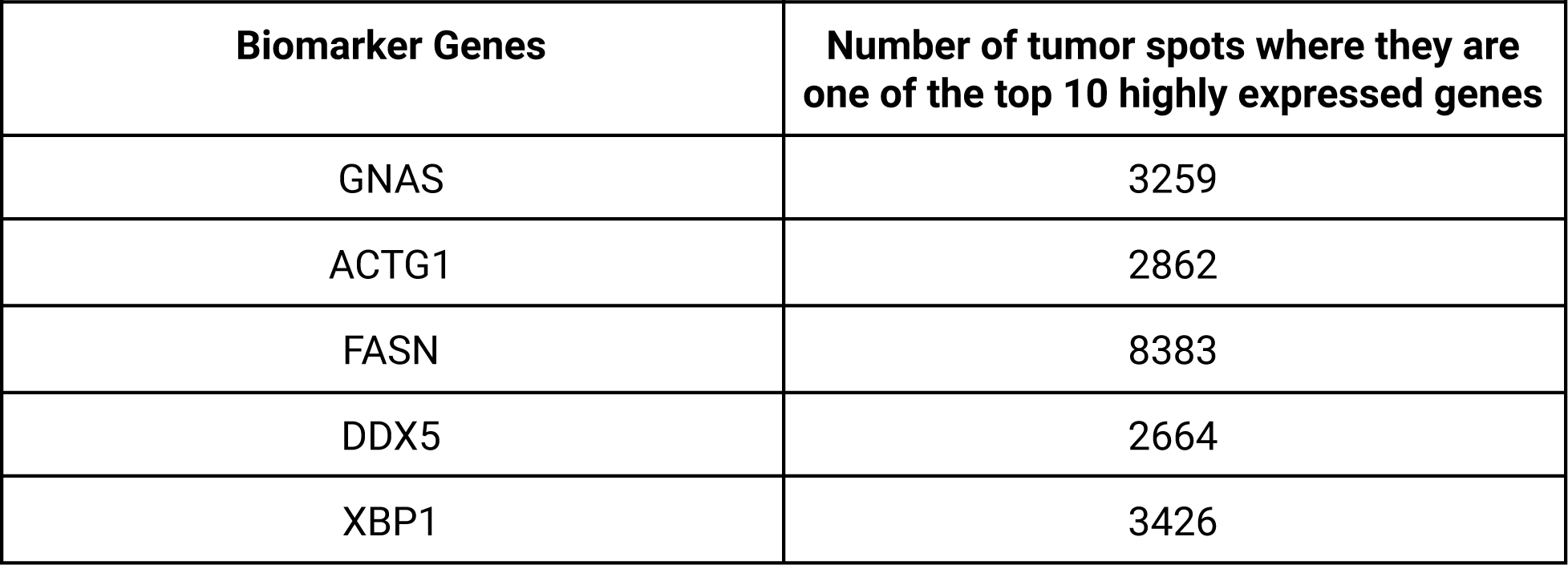
We enlist GNAS, ACTG1, FASN, DDX5 and XBP1 as 5 known biomarker genes from the gene list described in the 10x Visium breast cancer dataset. To observe the predicted expression of these genes across all the 30,612 tumor spots in this dataset; we sorted all the predicted gene expression values yielded by our co-trained model embeddings for each spot and picked the top 10 highly expressed genes across all the tumor spots to show the enrichment of these 5 known biomarker genes for breast cancer.

In the analysis shown in Figure 6., we compared the Spearman correlation coefficients for the predicted gene expression values against the actual gene expression values for the five chosen breast cancer biomarker genes (GNAS, ACTG1, FASN, DDX5, and XBP1) within the 10x Visium breast cancer dataset. We utilized four different types of spot representations or embeddings, namely :

a. CTransPath vision embeddings,
b. GIN graph embeddings,
c. Integrated CTransPath vision embeddings and GIN graph embeddings, and
d. Integrated embeddings co-trained on TCGA-BRCA and 10x Visium breast cancer tissue samples.

**Figure 6.**
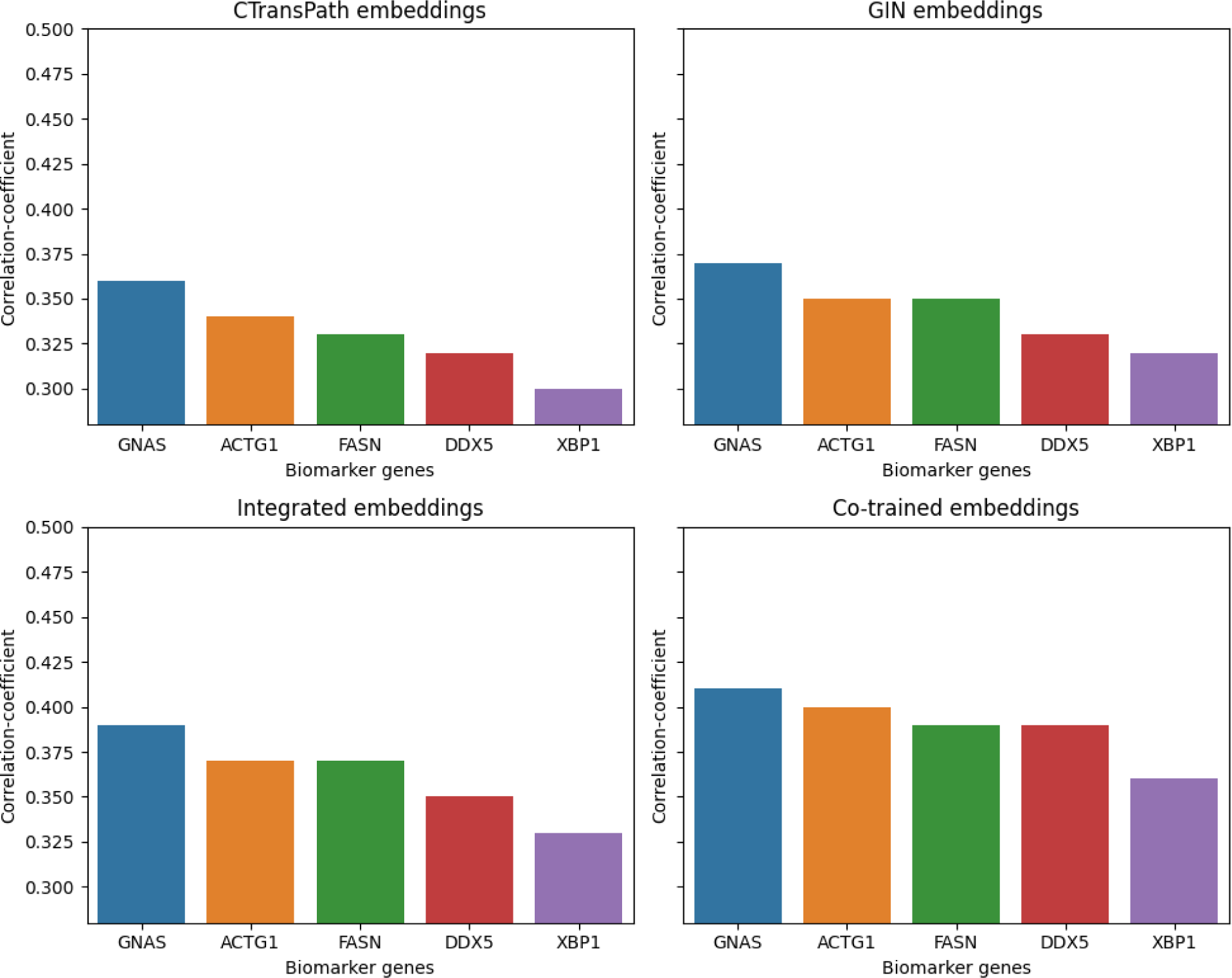
Comparison of the spearman correlation coefficient between the predicted and actual gene expression values for 5 breast cancer biomarker genes (GNAS, ACTG1, FASN, DDX5 and XBP1) in the 10x Visium breast cancer dataset using 4 different types of spot representations/embeddings : (a) CTransPath vision embeddings (b) GIN graph embeddings (c) Integrated CTransPath vision embeddings and GIN graph embeddings (d) Integrated embeddings co-trained on TCGA-BRCA and 10x Visium breast cancer tissue samples.

This analysis provides critical insights into the enrichment of known biomarker genes for breast cancer. Notably as shown in Figure 6 and Table 2, we observed that the correlation scores for all the biomarker genes were consistently higher when employing integrated embeddings, derived from both CTransPath vision and GIN graph embeddings, for spatial transcriptomic prediction, in contrast to using purely CTransPath vision embeddings or GIN graph embeddings. Most significantly, the co-trained embeddings exhibited the highest correlation scores for all five biomarker genes in the 10x Visium breast cancer dataset, outperforming the other three embedding types. This underscores the superior predictive power of co-trained embeddings in capturing the complex relationships governing gene expression within the spatial context of breast cancer tissue samples

**Table 2.** Table summarizing the spearman correlation coefficients for the predicted vs actual gene expression values for 5 breast cancer biomarker genes (GNAS, ACTG1, FASN, DDX5, XBP1) in the 10x Visium breast cancer dataset using 4 different types of spot representations/embeddings namely : CTransPath vision embeddings, GIN graph embeddings, Integrated CTransPath vision embeddings and GIN graph embeddings and Integrated embeddings co-trained on TCGA-BRCA and 10x Visium breast cancer tissue samples.

|  | **CTransPath** | **GIN** | **Integrated** | **Co-trained** |
| --- | --- | --- | --- | --- |
| **GNAS** | 0.36 | 0.37 | 0.39 | **0.41** |
| **ACTG1** | 0.34 | 0.35 | 0.37 | **0.40** |
| **FASN** | 0.33 | 0.35 | 0.37 | **0.39** |
| **DDX5** | 0.32 | 0.33 | 0.35 | **0.39** |
| **XBP1** | 0.30 | 0.32 | 0.33 | **0.36** |

We conducted comprehensive ablation studies to assess the individual and combined contributions of different components within our methodology. These ablation studies aimed to elucidate the impact of various strategies on the predictive performance of our model for spatial transcriptomic gene expression. We evaluated four different algorithmic scenarios, each focusing on specific aspects of our approach.

The first scenario involved utilizing only Graph Isomorphism Network (GIN) embeddings and regression for predicting spatial transcriptomic gene expression. This allowed us to understand the predictive power of GIN embeddings independently. The second scenario employed only CTransPath embeddings and regression to assess their standalone predictive capability. This analysis helped us evaluate the efficacy of CTransPath embeddings in capturing relevant spatial transcriptomic information. In the third scenario, we integrated both GIN and CTransPath embeddings, followed by regression. This joint approach was designed to leverage the strengths of both embedding methods to potentially enhance predictive performance. Finally, in the fourth scenario, we introduced co-training with TCGA BRCA bulk RNA Seq data, combining joint embeddings with regression. This additional step aimed to improve predictive power through cross-training with a different dataset.

To evaluate these scenarios, we measured Root Mean Squared Error (RMSE) and visualized gene expression spatialization for specific biomarker genes using heatmaps. The heatmaps allowed us to qualitatively assess how well the models captured the spatial patterns of gene expression across different algorithms. In Figure 7 for instance, we present a visual representation of the true and predicted expression patterns for the tumor biomarker GNAS within a patient sample using various spot representations or embeddings. The heatmaps depict the spatialization of GNAS gene expression across the tissue sample, offering valuable insights into its distribution. Remarkably, our observations reveal distinct differences among these representations. In the case of integrated embeddings and co-trained embeddings, denoted as (c) and (d), respectively, the resulting heatmaps closely align with the ground truth, effectively capturing the spatial distribution of GNAS. However, when employing CTransPath vision embeddings (a) or GIN graph embeddings (b), the spatialization of this biomarker gene appears less accurate. This observation, gleaned from Figure 7, underscores the advantages of utilizing integrated and co-trained embeddings for spatial transcriptomic predictions, as they outperform individual representations in capturing the spatial nuances of critical biomarker genes like GNAS.

**Figure 7.**
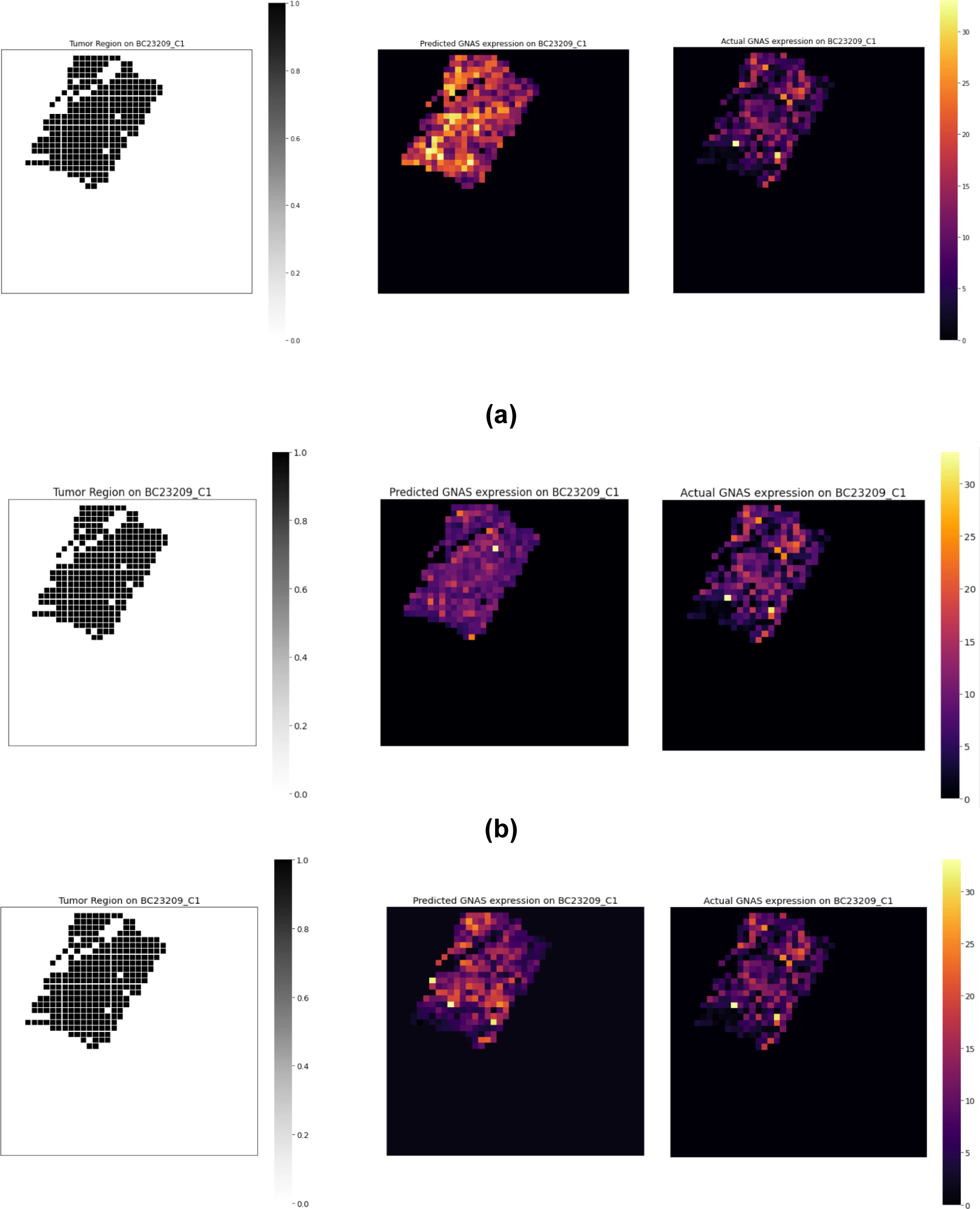
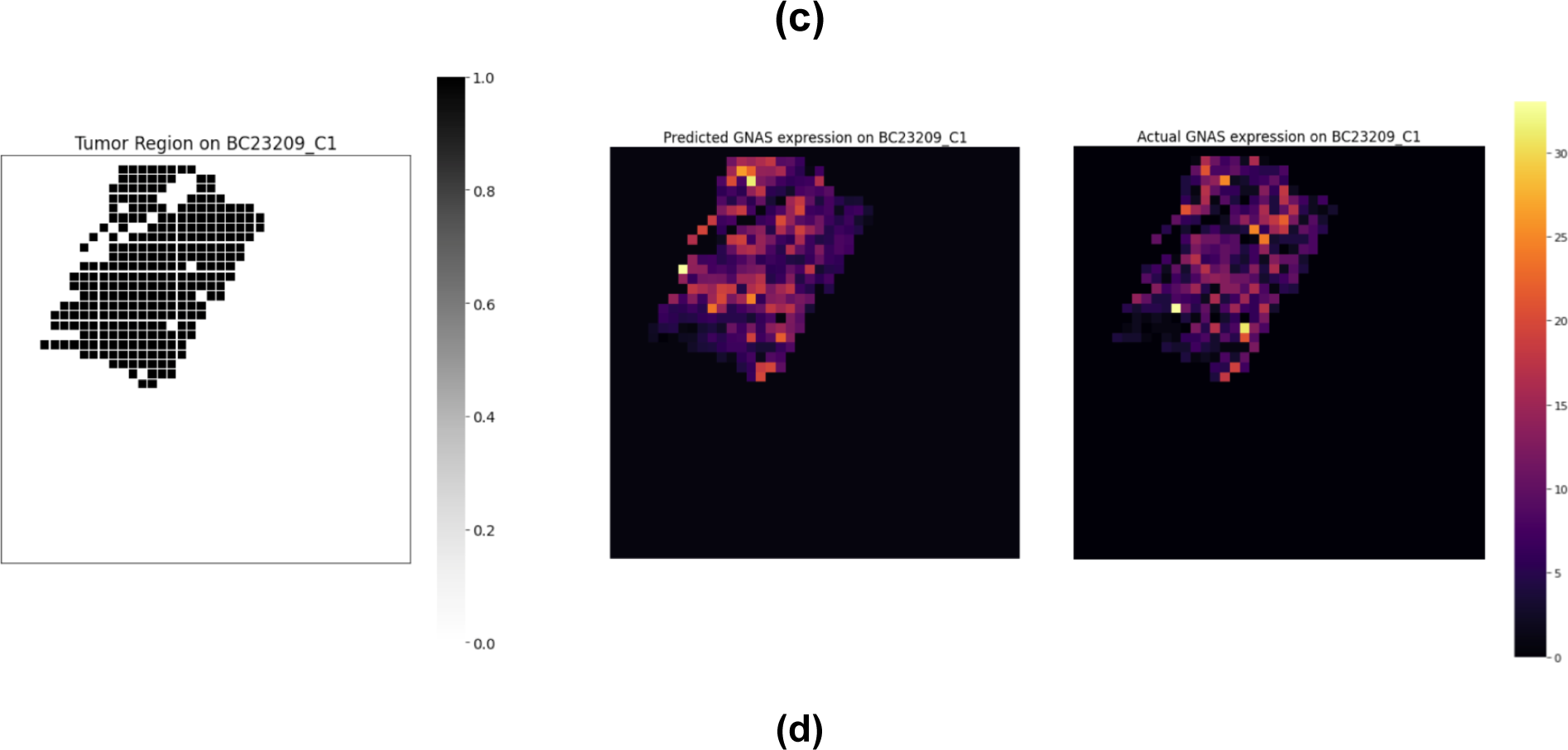
Visualization of the true and predicted expression for the tumor biomarker GNAS for a patient sample using (a) cTransPath vision embeddings (b) GIN graph embeddings (c) Integrated CTransPath vision embeddings and GIN graph embeddings (d) Integrated embeddings co-trained on TCGA-BRCA and 10x Visium breast cancer tissue samples.

We also examined the efficacy of the four distinct spot representations or embeddings in the context of downstream multioutput regression for predicting spatial transcriptomics. Employing a leave-one-patient-out cross-validation strategy, we assessed the root mean square error (RMSE) across each validation fold, subsequently computing an average RMSE as a performance metric. Notably, our findings, as depicted in Table 3, revealed a distinct hierarchy of performance among these embeddings. Co-trained embeddings exhibited the lowest cross-validated RMSE, highlighting their superior predictive power. Following closely were the integrated embeddings, which also delivered competitive results. Surprisingly, GIN graph embeddings and CTransPath vision embeddings yielded quite similar cross-validated RMSE values, suggesting comparable performance in the context of downstream regression. This observation underscores the significant advantage of co-trained and integrated embeddings in enhancing the predictive accuracy of spatial transcriptomics compared to their individual counterparts.

**Table 3.** Table summarizing the cross-validated RMSE for downstream multi-output regression task to predict spatial transcriptomics across all 23 patient profiles in the 10x Visium breast cancer dataset using leave one patient out cross-validation setup across 4 different types of spot representations/embeddings namely: CTransPath vision embeddings, GIN graph embeddings, Integrated CTransPath vision embeddings and GIN graph embeddings and Integrated embeddings co-trained on TCGA-BRCA and 10x Visium breast cancer tissue samples.

|  | **CTransPath**<br>**Embeddings** | **GIN**<br>**Embeddings** | **Integrated**<br>**Embeddings** | **Co-trained**<br>**Embeddings** |
| --- | --- | --- | --- | --- |
| **Cross-validated**<br>**RMSE** | 0.33 ± 0.038 | 0.32 ± 0.039 | 0.27 ± 0.035 | **0.24 ± 0.028** |

Our ablation studies provided valuable insights into the contributions of individual components and their synergistic effects, shedding light on the most effective strategies for spatial transcriptomic gene expression prediction.

## Interpretability of results

Interpreting the results of our co-training model is crucial for gaining insights into the underlying biological mechanisms. To achieve this, we employed a multi-step interpretability strategy as described in Figure 8. First, we identified spot representations (embeddings) for the 10x Visium tiles encompassing these spots and clustered these spot representations/tile embeddings, along with the tile embeddings from the TCGA BRCA dataset. By examining the clusters formed, we could observe similarities and differences in the spatial organization of gene expression profiles between the two datasets. To delve deeper, we applied an instance of the GNNExplainer algorithm over 10x Visium and TCGA-BRCA tiles that were allocated to the same cluster [36]. This allowed us to rank the importance of nodes and edges within the graph corresponding to each tile in the cluster. This provided us with valuable information on the specific cell types and cell-cell interactions contributing to the model’s predictions. By running GNNExplainer on both the 10x Visium and TCGA tiles for the spatial transcriptomic prediction task, we could identify and compare the nodes/cell types responsible for the maximum gradient change. This comprehensive interpretability approach enhances our understanding of the biological relevance of our model’s predictions and highlights key factors influencing gene expression patterns in histopathological samples.

**Figure 8.**
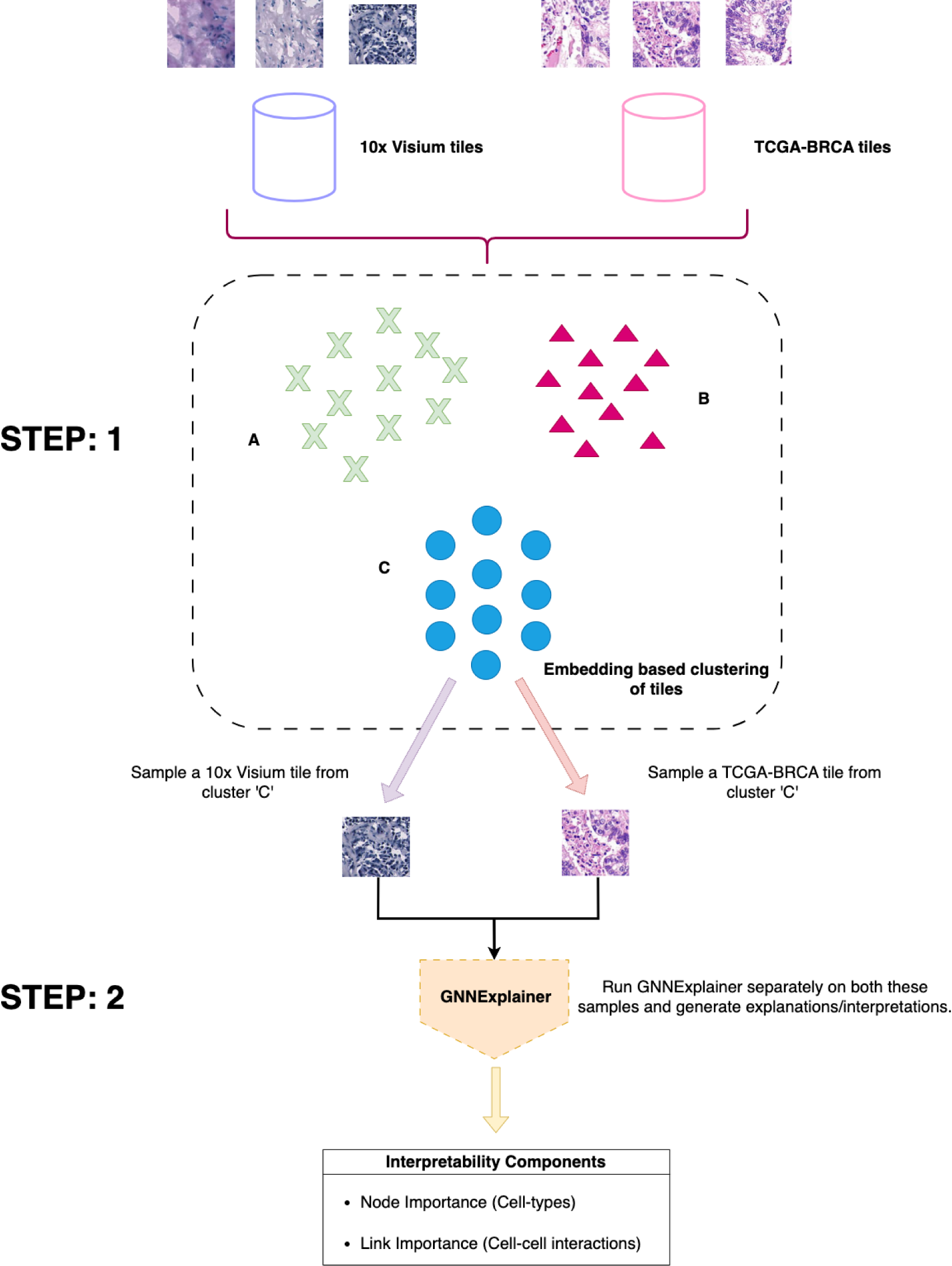
Overview of our multi-step strategy for producing cellular explanations/interpretations in the context of spatial transcriptomics prediction using the co-trained model embeddings.

### Interpretation of our model’s insights from 10x Visium breast cancer dataset

We carefully selected a breast cancer tissue sample, denoted as BC23209 C1, from the 10x Visium breast cancer dataset. Additionally, we identified a corresponding sample from the TCGA-BRCA dataset, specifically labeled as TCGA-3C-AAAU-01A-01-TS1. It’s worth highlighting that both of these samples were grouped into the same cluster during our initial embedding-based clustering process. Subsequently, we conducted GNNExplainer analysis on these two samples, aiming to explore the commonalities that led to their clustering together, primarily focusing on their embedding similarities.

The GNNExplainer scoring shown in Table 4(a) and Table 4(b) highlighted four key cell types of significant importance: Lymphocytes, Epithelial cells, Macrophages, and Neutrophils. Notably, Lymphocytes emerged as a central player, showing strong influence over Macrophages and Neutrophils, thereby signifying their integral role in orchestrating immune responses within the breast cancer tissue. Epithelial cells, forming the foundational structure of the breast tissue, held their own significance, mediating interactions with both Lymphocytes and Macrophages. This insightful ranking underscores the complex interplay of these cell types within the tumor microenvironment, shedding light on their critical contributions to spatial transcriptomic patterns.

**Table 4.** (a) Aggregated node importance scores for each cell-type from the GNNExplainer on the 10x Visium sample. (b) Aggregated link importance scores for each cell-cell interaction from the GNNExplainer on the 10x Visium sample.

| Cell-type/Node | Aggregated node importance score from GNNExplainer |
| --- | --- |
| Lymphocytes | 0.88 |
| Epithelial cell | 0.82 |
| Macrophages | 0.68 |
| Neutrophils | 0.61 |

| Cell-cell interactions | Aggregated link importance score from GNNExplainer |
| --- | --- |
| Epithelial cell - Lymphocytes | 0.91 |
| Lymphocytes - Macrophages | 0.87 |
| Epithelial cell - Macrophages | 0.82 |
| Lymphocytes - Neutrophils | 0.77 |
| Epithelial cell - Neutrophils | 0.69 |
| Macrophages - Neutrophils | 0.62 |

### Interpretation of our model’s insights from TCGA-BRCA dataset

Under the same experimental conditions and samples utilized when delineating the interpretation of our model’s insights from 10x Visium breast cancer dataset; we repeated the interpretability experiment and extracted insights from GNNExplainer on the TCGA-BRCA sample - TCGA-3C-AAAU-01A-01-TS1.

As shown in Table 5(a) and Table 5(b), the GNNExplainer analysis conducted on the TCGA-BRCA sample spotlighted the prominent roles of Lymphocytes, Epithelial cells, Neutrophils, and Macrophages, with each of them contributing distinctly to the gradient changes.

**Table 5.** (a) Aggregated node importance scores for each cell-type from the GNNExplainer on the TCGA-BRCA sample. (b) Aggregated link importance scores for each cell-cell interaction from the GNNExplainer on the TCGA-BRCA sample.

| Cell-type/Node | Aggregated node importance score from GNNExplainer |
| --- | --- |
| Lymphocytes | 0.91 |
| Epithelial cell | 0.79 |
| Neutrophils | 0.63 |
| Macrophages | 0.57 |

| Cell-cell interactions | Aggregated link importance score from GNNExplainer |
| --- | --- |
| Epithelial cell - Lymphocytes | 0.85 |
| Lymphocytes - Macrophages | 0.82 |
| Epithelial cell - Macrophages | 0.76 |
| Epithelial cell - Neutrophils | 0.66 |
| Macrophages - Neutrophils | 0.64 |
| Lymphocytes - Neutrophils | 0.60 |

Upon closer examination of the link importance, we observed intricate relationships within these cell types. Epithelial cells held sway over Lymphocytes, Macrophages, and Neutrophils, suggesting their pivotal role as a regulatory hub. Lymphocytes, in turn, influenced Macrophages, fostering immune responses within the tissue. Additionally, Macrophages exhibited influence over Neutrophils, hinting at their interconnected roles in the context of immune response and tissue regulation.

Comparing this analysis with our previous study on the 10x Visium dataset, we found remarkable similarities. In both cases, Lymphocytes, Epithelial cells, Macrophages, and Neutrophils emerged as key players, emphasizing their significance in orchestrating spatial transcriptomics. This alignment underscores the robustness and generalizability of our findings across different datasets, enhancing our understanding of the complex interplay between gene expression and tissue architecture in breast cancer.

## Conclusion and Future work

In conclusion, our research has demonstrated the effectiveness of a novel approach for spatial transcriptomic gene expression prediction, leveraging Graph Isomorphism Network (GIN) embeddings, CTransPath embeddings, and co-training with TCGA BRCA bulk RNA Seq data. Our comprehensive evaluations, utilizing Mean Squared Error (MSE), Spearman correlation coefficients, and AUROC, highlighted the robustness and predictive power of our model across different datasets and tasks. By co-training with TCGA BRCA data, we achieved improved performance for both spatial transcriptomic gene expression and whole slide image-level bulk RNA Seq predictions. Our findings underscore the potential of combining deep learning strategies with diverse data sources to enhance our understanding of complex biological phenomena.

Future work in this direction could focus on further refining the integration of spatial transcriptomic data and bulk RNA Seq data, exploring additional self-supervised learning strategies, and extending our methodology to other cancer types and diseases. Additionally, incorporating more comprehensive biological annotations and domain-specific knowledge could enhance the interpretability and clinical relevance of our predictions. Moreover, the development of user-friendly tools and platforms to facilitate the adoption of our approach by the broader scientific community would be valuable for accelerating research in the field of spatial transcriptomics and histopathological analysis.

## Acknowledgements

We would like to thank all of the study participants and their families, and all of the site investigators, study coordinators and staff. This work was supported by Genentech, Inc.

## Data Availability

You can access all H&E whole slide images and processed data by visiting http://www.spatialtranscriptomicsresearch.org. If you’re interested in the 10x Spatial Genomics data then you can download the corresponding H&E whole slide images and relevant spatial transcriptomic ground truths for the spots within each H&E whole slide image from https://wp.10xgenomics.com/spatial-transcriptomics. Additionally, all TGCA data is publicly accessible through the Genomic Data Commons Data Portal at https://portal.gdc.cancer.gov.

## Author contributions

K.L.,V.P. devised the study.

E.W., V.P. preprocessed and analyzed the data.

V.P. wrote the manuscript with input from the remaining authors.

## Competing interests

K.L. is an employee of Genentech, Inc. and shareholder in F. Hoffmann La Roche, Ltd.

V.P. is an external partner at Genentech, Inc.

E.W. was an intern at Genentech, Inc.

